# Intra-condensate demixing of TDP-43 inside stress granules generates pathological aggregates

**DOI:** 10.1101/2024.01.23.576837

**Authors:** Xiao Yan, David Kuster, Priyesh Mohanty, Jik Nijssen, Karina Pombo-García, Azamat Rizuan, Titus M. Franzmann, Aleksandra Sergeeva, Patricia M. Passos, Leah George, Szu-Huan Wang, Jayakrishna Shenoy, Helen L. Danielson, Alf Honigmann, Yuna M. Ayala, Nicolas L. Fawzi, Jeetain Mittal, Simon Alberti, Anthony A. Hyman

## Abstract

Cytosolic aggregation of the nuclear protein TDP-43 is associated with many neurodegenerative diseases, but the triggers for TDP-43 aggregation are still debated. Here, we demonstrate that TDP-43 aggregation requires a double event. One is up-concentration in stress granules beyond a threshold, and the other is oxidative stress. These two events collectively induce intra-condensate demixing, giving rise to a dynamic TDP-43 enriched phase within stress granules, which subsequently transitions into pathological aggregates. Mechanistically, intra-condensate demixing is triggered by local unfolding of the RRM1 domain for intermolecular disulfide bond formation and by increased hydrophobic patch interactions in the C-terminal domain. By engineering TDP-43 variants resistant to intra-condensate demixing, we successfully eliminate pathological TDP-43 aggregates in cells. We conclude that up-concentration inside condensates and simultaneous exposure to environmental stress could be a general pathway for protein aggregation, with intra-condensate demixing constituting a key intermediate step.

## INTRODUCTION

A key hallmark of most neurodegenerative diseases is the formation of protein aggregates. For instance, aggregates of Tau protein are associated with Alzheimer’s disease, and aggregates of α-synuclein are associated with Parkinson’s disease. Test tube experiments with purified disease-causing proteins have shown that proteins in isolation are soluble in solution.^1,2^ Their aggregation typically is a rare, concentration-dependent event that requires a structural destabilization induced by changes in physical-chemical conditions or genetic mutations. In cells the very same proteins are typically highly soluble, despite the crowded intracellular milieu.^3^ For example, overexpression of Tau or α-synuclein is insufficient to generate protein aggregates in cells.^4–7^

The stability of disease-causing proteins poses the following question: Where and how do pathological protein aggregates arise in the intracellular environment? In particular, what are the actual triggers for a soluble protein to become insoluble and initiate the aggregation process? One of the most devastating neurodegenerative diseases is amyotrophic lateral sclerosis (ALS), a motor neuron disease that commonly results in loss of motor neurons and death within a few years of onset of symptoms. The protein most frequently associated with ALS is the nuclear TAR DNA-binding protein 43 (TDP-43). TDP-43 has essential functions in transcriptional regulation, pre-mRNA splicing and translation regulation.^8–11^ TDP-43 predominantly localizes to the nucleus because of a nuclear localization signal that is recognized by importins.^12,13^ The cytosolic TDP-43 concentration is low under physiological conditions, but under stress it passively leaks into the cytoplasm.^14^ Mis-localized cytosolic TDP-43 forms aggregates in ∼97% of ALS, ∼45% of frontotemporal dementia (FTD), and all cases of Alzheimer’s disease-associated LATE (Limbic-predominant Age-related TDP-43 Encephalopathy), suggesting that nuclear depletion and cytoplasmic aggregation of TDP-43 is a general hallmark of several age-related neurodegenerative diseases.^15–19^ The cytoplasmic accumulation and aggregation of TDP-43 could confer a loss-of-function by mis-localizing, and/or a gain-of-toxicity by sequestration of essential cellular factors.^13,20–24^ Therefore, a central challenge in ALS research is to identify the triggers for wild-type TDP-43 aggregation.

The aggregation of TDP-43 is thought to be driven by a structural change of an α-helical region in the C-terminus that transitions into a cross-β sheet structure.^25,26^ This structural transition has been shown to be accelerated by disease mutations in the α-helical region,^27^ although TDP-43 aggregation also occurs in the absence of mutations in most ALS cases. However, although the structural transitions that lead to aggregated TDP-43 are well studied, the actual triggers that induce the onset of aggregation in the cellular context remain unknown. Oxidative stress has been implicated as an environmental trigger,^28^ but the relationship between oxidative stress and structural destabilization of TDP-43 remains to be determined. This lack of understanding hinders the development of therapeutic agents.

Human genetics suggests a strong connection between ALS and a type of biomolecular condensate called a stress granule.^29–33^ Stress granules are canonical condensates in the eukaryotic cytosol that are triggered by various stresses and assemble *via* interactions between RNA and RNA-binding proteins.^34–36^ Because many of these stress granule-associated RNA-binding proteins have been linked to ALS, it has been hypothesized that stress granules could in part be crucibles of such diseases.^31^ However, the role of cellular condensates in promoting aggregation of TDP-43 is controversial. Patient data have shown that pathological TDP-43 aggregates contain key stress granule proteins such as TIA-1, eIF3, and PABP-1.^37–39^ Indeed, a recent histopathological examination, using patient samples, indicated the presence of the stress granule marker HuR in early-stage ALS spinal cord inclusions.^40^ However, other work has suggested that aggregation occurs outside rather than inside stress granules and has proposed that stress granules might protect against aggregation.^24,41–46^ These disparate results have made it difficult to come to any general conclusions about the role of stress granule condensates in TDP-43 aggregation.

Here, we investigate the role of stress granules in TDP-43 aggregation. Using biochemical reconstitution, biophysical analysis, computational simulations and genetic experiments in cells, we show that aggregation of TDP-43 requires a double event. One event is that stress granules raise the concentration of cytosolic TDP-43 above a critical threshold. The other event is oxidative stress, which leads to cysteine oxidation. Together, these two events induce an intra-condensate demixing process, giving rise to a TDP-43 enriched phase within stress granules. The demixing process is driven by local unfolding of the RRM1 domain for cysteine oxidation and disulfide bond formation, and by increased hydrophobic patch interactions in the C-terminal domain. These condensates then harden to form aggregates with pathological hallmarks, in both HeLa cells but also neurons derived from iPS cells. By engineering TDP-43 variants that are resistant to intra-condensate demixing, we successfully eliminate pathological TDP-43 aggregates in cells. Together, our findings reveal that stress granules can act as crucibles triggering TDP-43 aggregation *via* a key intra-condensate demixing process. We suggest the combination of increased concentration within condensates and simultaneous exposure to environmental stress could be a general mechanism for *de novo* protein aggregation in cells.

## RESULTS

### TDP-43 undergoes intra-condensate demixing inside stress granules

TDP-43 is primarily a nuclear protein under physiological conditions, where it regulates the splicing of multiple genes. Cytoplasmic aggregation requires it to leave the nucleus. To decouple the process of nucleocytoplasmic shuttling from aggregation in the cytoplasm, we transfected HeLa cells with a GFP-tagged TDP-43 variant in which the nuclear localization signal (NLS) was disrupted (TDP-43^ΔNLS^).^47^ After stressing cells by adding 100 μM arsenite, we noticed three different cellular phenotypes. Two of those have previously been observed: 1) cells with dispersed TDP-43^ΔNLS^ in stress granules,^37^ 2) cells exhibiting large TDP-43^ΔNLS^ aggregates independent of stress granules.^24,46^ However, we also noticed that in many cells TDP-43^ΔNLS^ showed a punctate distribution inside stress granules.

Plasmid transfection resulted in populations of cells with a ten-fold variation in cytosolic TDP-43^ΔNLS^ concentration. To investigate whether the observed phenotypes are related to protein concentration, we determined cytosolic TDP-43^ΔNLS^ expression levels before stress (**Figures 1A** and **S1A**). Cells exhibiting the above three distinct behaviors correlated strongly with low (∼0.2–0.4 μM), high (∼0.8–4.0 μM) and medium (∼0.4–0.7 μM) expression of TDP-43^ΔNLS^ (**Figure 1A**). It is noteworthy that the endogenous concentration of TDP-43 in HeLa cells is approximately 0.5 μM,^48^ which falls within the medium expression range but not in high expression cells.

**Figure 1.**
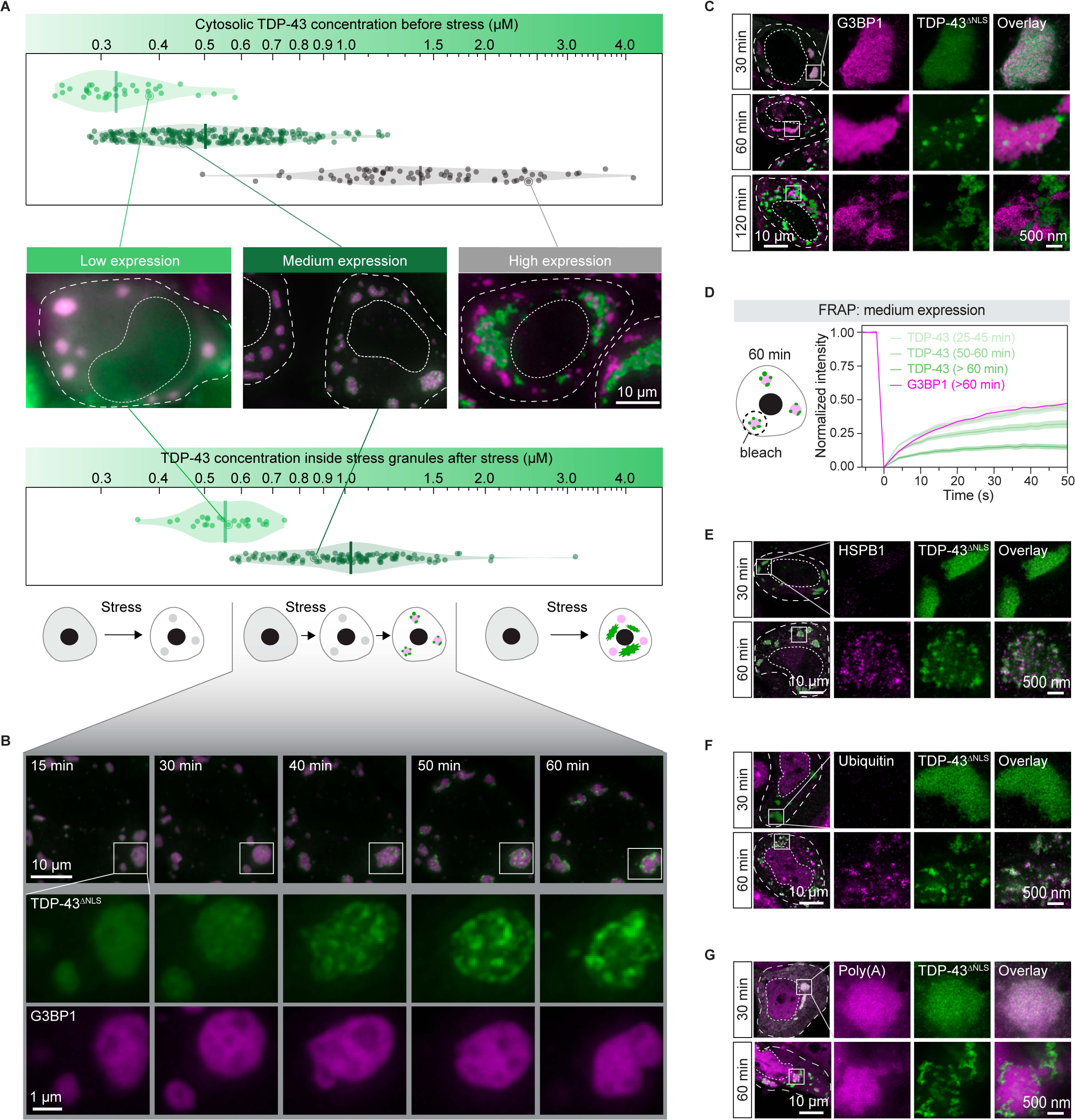
TDP-43 undergoes intra-condensate demixing inside stress granules and results in pathological aggregates (A) Three different cellular phenotypes of TDP-43 in HeLa cells. The cytosolic concentration of TDP-43^ΔNLS^ in low, medium and high expression cells was measured before stress (top), and TDP-43^ΔNLS^ concentration inside mCherry-tagged G3BP1 stress granules was measured right before its intra-condensate demixing under stress with 100 μM arsenite (bottom). The cellular and nucleus boundaries are indicated by dashed lines (middle). Scale bar, 10 μm. (B) The time evolution of a representative stress granule with intra-condensate demixing of TDP-43^ΔNLS^ in medium expression cells. Stress granules marked by squares under confocal fluorescence microscopy were zoomed in. Scale bar, 10 μm and 1 μm for confocal and zoomed in images, respectively. (C) Representative STED images of HeLa cells expressing TDP-43^ΔNLS^ after addition of 100 μM arsenite from 30 min to 120 min. Stress granules marked by squares under confocal microscopy were further visualized by STED microscopy. Scale bar, 10 μm and 500 nm for confocal and STED images, respectively. (D) FRAP of TDP-43^ΔNLS^ and mCherry-tagged G3BP1 in medium expression cells after addition of 100 μM arsenite over time. Data represent the mean ± SD. (E) HSPB1 recognition of TDP-43 aggregates upon intra-condensate demixing by STED. Cells expressing TDP-43^ΔNLS^ were stressed with 100 μM arsenite and images were acquired before and after demixing. Scale bar, 10 μm and 500 nm for confocal and STED images, respectively. (F) Ubiquitination of TDP-43 aggregates upon intra-condensate demixing by STED. Cells expressing TDP-43^ΔNLS^ were stressed with 100 μM arsenite and images were acquired before and after demixing. Scale bar, 10 μm and 500 nm for confocal and STED images, respectively. (G) mRNA-FISH assay in HeLa cells expressing TDP-43^ΔNLS^ by STED. Cells expressing TDP-43^ΔNLS^ were stressed with 100 μM arsenite. Atto647N-labeled oligo-dT oligonucleotides were used to stain poly(A)-containing mRNA. Images were acquired before and after demixing. Scale bar, 10 μm and 500 nm for confocal and STED images, respectively. See also Figure S1 and Videos S1–S3.

In low expression cells, TDP-43^ΔNLS^ was diffuse in the cytoplasm before stress (**Figure S1**), and after stress accumulated in stress granules where it remained dispersed and dynamic, as assayed by fluorescence recovery after photobleaching (FRAP) (**Figures 1A**, **S1C**, and **Video S1**). In high expression cells, TDP-43^ΔNLS^ assembled into condensates before stress, which were dynamic, with an average diameter of ∼1.1 μm (**Figures S1B** and **S1D**). Upon adding 100 μM arsenite to stress the cells, these TDP-43^ΔNLS^ condensates rapidly transformed into aggregates (**Figure S1D**), and remained independent of stress granules (**Figure 1A** and **Video S2**). This transition of dynamic condensates to less dynamic aggregates is in agreement with previous studies.^24,46^

In medium expression cells, TDP-43^ΔNLS^ was diffuse before stress (**Figure S1B**). However, after stress, TDP-43^ΔNLS^ was locally concentrated and formed puncta inside stress granules (**Figure 1A**). The time evolution of a representative stress granule exhibiting TDP-43^ΔNLS^ puncta is shown in **Figure 1B** and **Video S3**. Initially, TDP-43^ΔNLS^ was uniformly dispersed inside stress granules, but after 30 min of stress, distinct TDP-43 puncta emerged that excluded the stress granule marker G3BP1 (**Figure 1B**). We name this phenomenon of TDP-43^ΔNLS^ puncta formation inside stress granules *intra-condensate demixing*, as it is accompanied by the formation of two distinct condensed phases.

Stimulated emission depletion microscopy imaging (STED; see STAR Methods) confirmed that intra-condensate demixing occurs (**Figure 1C**). With longer exposure to stress (120 min), TDP-43^ΔNLS^ showed marked separation from G3BP1. This was accompanied by a change in TDP-43^ΔNLS^ morphology from the distributed spherical condensates to irregularly shaped structures, eventually giving rise to fragmented stress granules (**Figure 1C**). We also confirmed that intra-condensate demixing occurs for wild-type TDP-43 containing a functional NLS (**Figure S1E**).

### Intra-condensate demixing of TDP-43 results in pathological aggregation

Demixed TDP-43 puncta exhibited a liquid-like behavior, as indicated by droplet fusion (**Figure S1F**). However, the dynamics of TDP-43 protein gradually slowed over time, as studied by FRAP analysis (**Figure 1D**). The internal concentration of individual TDP-43 puncta was further elevated compared to their state before demixing inside stress granules (**Figure S1G**), a process that has been coupled to a liquid-to-solid transition.^49^

We tested whether the demixed TDP-43 shows aggregation hallmarks using three commonly used markers: recruitment of heat shock proteins, ubiquitination and phosphorylation. Ubiquitination and phosphorylation are two commonly found pathological hallmarks of TDP-43 inclusions in ALS and FTD patients.^15,50,51^ HSPB1, a small heat shock protein which was shown previously to interact with misfolded TDP-43,^46^ weakly localized to stress granules before intra-condensate demixing (**Figure 1E**). However, upon prolonged stress, HSPB1 strongly co-stained with demixed TDP-43^ΔNLS^ puncta (**Figure 1E**). TDP-43^ΔNLS^ puncta also exhibited strong ubiquitin signal and were recognized with an antibody specific for phosphorylation at Ser409/410 (**Figures 1F** and **S1H**).

By visualizing poly(A)-containing mRNA using an Atto647N-labeled oligo-dT oligonucleotide, we observed that the colocalization between demixed TDP-43^ΔNLS^ and mRNA was lost upon prolonged stress (**Figure 1G**). This suggests that TDP-43 is losing its structure or function as an RNA-binding protein.^11,52^ Demixed TDP-43^ΔNLS^ puncta recruited nuclear TDP-43 into these cytoplasmic aggregates (**Figure S1I**), suggesting that aggregates generated *via* intra-condensate demixing could cause a gain-of-toxicity effect by sequestering key cellular factors.^33,53,54^

Taken together, our data so far show that if the TDP-43 concentration exceeds a threshold concentration after partitioning into stress granules, it will undergo intra-condensate demixing and assemble into a TDP-43 enriched phase. Inside this phase, TDP-43 dynamics slow down and the protein starts to form pathological aggregates.

### Intra-condensate demixing of TDP-43 promotes a liquid-to-solid phase transition in reconstituted stress granules

We used *in vitro* reconstitution to demonstrate that intra-condensate demixing is a distinct phase separation-driven process within multicomponent stress granules. For this purpose, we employed a minimal stress granule system that is based on purified G3BP1 and RNA^34^ (**Figure 2A**; see STAR Methods). TDP-43 readily partitioned into these minimal stress granules and colocalized well with G3BP1 (**Figure 2A**). Strikingly, initially well-mixed TDP-43 formed a separate phase inside stress granules over time, which excluded G3BP1 (**Figure 2B** and **Video S4**), similar to the intra-condensate demixing observed in cells (**Figure 1B**). By calculating the Pearson correlation between the fluorescence intensities of TDP-43 and G3BP1 and setting an apparent demixing threshold of 0.85, TDP-43 demixing occurred after 5–6 h (**Figure 2B**). Quantifying TDP-43 concentration before demixing (0–4 h) revealed that stress granules continuously concentrated TDP-43 to cross a threshold for intra-condensate demixing (**Figure 2C**).

**Figure 2.**
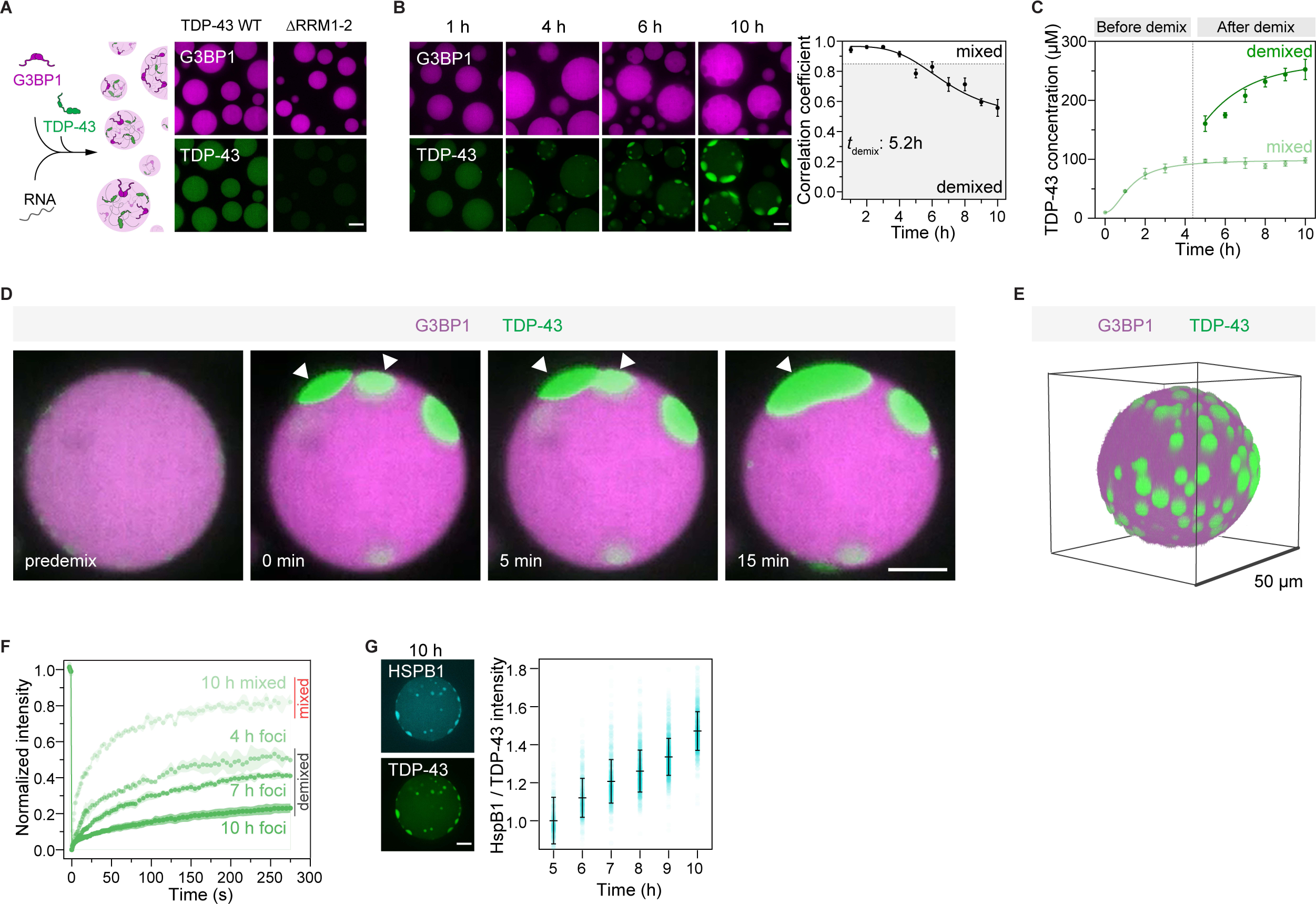
Intra-condensate demixing of TDP-43 promotes a liquid-to-solid phase transition in reconstituted stress granules (A) TDP-43 recruitment into minimal stress granules. Recombinant G3BP1 (20 μM) and Poly(A) RNA (80 ng/μl) were incubated to form minimal stress granules. TDP-43 WT or ΔRRM1-2 (0.5 μM) was included as a client of stress granules. Scale bar for Figure 2, 10 μm. (B) Intra-condensate demixing of TDP-43 inside minimal stress granules. TDP-43 (10 μM) was added into minimal stress granules in the presence of 2.5% dextran. Pearson colocalization between G3BP1 and TDP-43 was analyzed by Coloc 2 plugin in Fiji, and 0.85 was set as an apparent demixing threshold. Data represent the mean ± SD. (C) TDP-43 concentrations in the mixed and demixed phases within minimal stress granules along the intra-condensate demixing process. The time point for demixing is indicated by the dash line. (D) Representative images of demixed TDP-43 puncta fusion. Puncta undergoing fusion are indicated by white arrowheads. (E) 3D reconstruction of minimal stress granules upon intra-condensate demixing of TDP-43 at 10 h. (F) FRAP of TDP-43 in the mixed and demixed phases inside minimal stress granules along the intra-condensate demixing process. Data represent the mean ± SD. (G) Increasing recognition of demixed TDP-43 by HSPB1 along the intra-condensate demixing process. Alexa 546-labeled HSPB1 (0.5 μM) was added into minimal stress granules and the ratio of fluorescent intensity between HSPB1 and TDP-43 was measured to show the increased stoichiometry and represented as mean ± SD. Representative images of HSPB1 and TDP-43 at 10 h are shown. See also Figure S2 and Video S4.

Deleting the RNA-recognition motifs RRM1 and RRM2 (ΔRRM1-2) strongly reduced TDP-43 partitioning into stress granules (**Figure 2A**), suggesting that TDP-43 recruitment depends on its RNA binding. However, when demixed, the TDP-43-rich phase contained less RNA than the G3BP1-rich phase (**Figure S2A**), consistent with loss of RNA binding in cells (**Figure 1G**). Demixed TDP-43 puncta fused (**Figure 2D**), suggesting their liquid-like property. TDP-43 puncta tended to orient towards the periphery of stress granules (**Figure 2E**). Thus, our *in vitro* system showed that reconstituted minimal stress granules can drive the formation of a TDP-43 enriched phase, giving rise to multiphasic condensates.

After intra-condensate demixing, TDP-43 concentration remained constant in the mixed phase, while it progressively increased in the demixed phase (5–10 h, **Figure 2C**), indicating that demixed TDP-43 phase may undergo a liquid-to-solid transition.^49^ Evidence that the demixed TDP-43 phase leads to TDP-43 misfolding and aggregation came from the following experiments: First, mixed TDP-43 inside stress granules was dynamic by FRAP analysis, while demixed TDP-43 displayed aging over time and protein dynamics gradually slowed down (**Figures 2F** and **S2B**). Second, the aging of demixed TDP-43 also coincided with increased staining with an aggregation-specific amytracker dye and detection of oligomers by semi-denaturing agarose gel electrophoresis (SDD-AGE) (**Figures S2C** and **S2D**). Third, HSPB1 specifically recognized demixed TDP-43, and could counteract TDP-43 demixing in a concentration-dependent manner (**Figures 2G** and **S2E**). In agreement with the cellular experiments, TDP-43 misfolds and forms aggregates in the demixed phase, which can be prevented by chaperones.

Intra-condensate demixing of TDP-43 also occurred inside lysate stress granules reconstituted by adding recombinant G3BP1 to cellular extract^55^ (**Figures S2F–S2H**; see STAR Methods). Therefore, intra-condensate demixing of TDP-43 was observed in three distinct environments: 1) a minimal reconstituted system, 2) cellular extract, and 3) cells. In all cases, TDP-43 partitions into stress granules first, where it then demixes into a TDP-43 enriched phase. The TDP-43 phase undergoes a liquid-to-solid transition and eventually converts into protein aggregates.

### Oxidation is a prerequisite for intra-condensate demixing of TDP-43

We next investigated whether environmental insult that triggers protein misfolding is also required for intra-condensate demixing. In our cellular experiments, we used arsenite stress to trigger stress granule assembly, which most likely acts by generating reactive oxygen species (ROS) and causing oxidative stress.^56^ Oxidative stress has been implicated in the progression of ALS by triggering TDP-43 aggregation.^28,57,58^

We treated cells with different stressors. The milder oxidant paraquat also induced TDP-43^ΔNLS^ demixing like arsenite (**Figure S3A**). However, when stress granules were induced without oxidative stress by using puromycin,^59^ TDP-43^ΔNLS^ did not undergo intra-condensate demixing and remained dynamic inside stress granules, even at higher TDP-43 concentrations (**Figures 3A**, **3B** and **S3B**). Additionally, disturbing the protein quality control machinery by inhibiting proteasome or Hsp70 did not trigger TDP-43^ΔNLS^ demixing in the puromycin-induced stress granules (**Figure S3A**). Thus, oxidative stress is required for TDP-43 demixing and the subsequent generation of aggregates.

**Figure 3.**
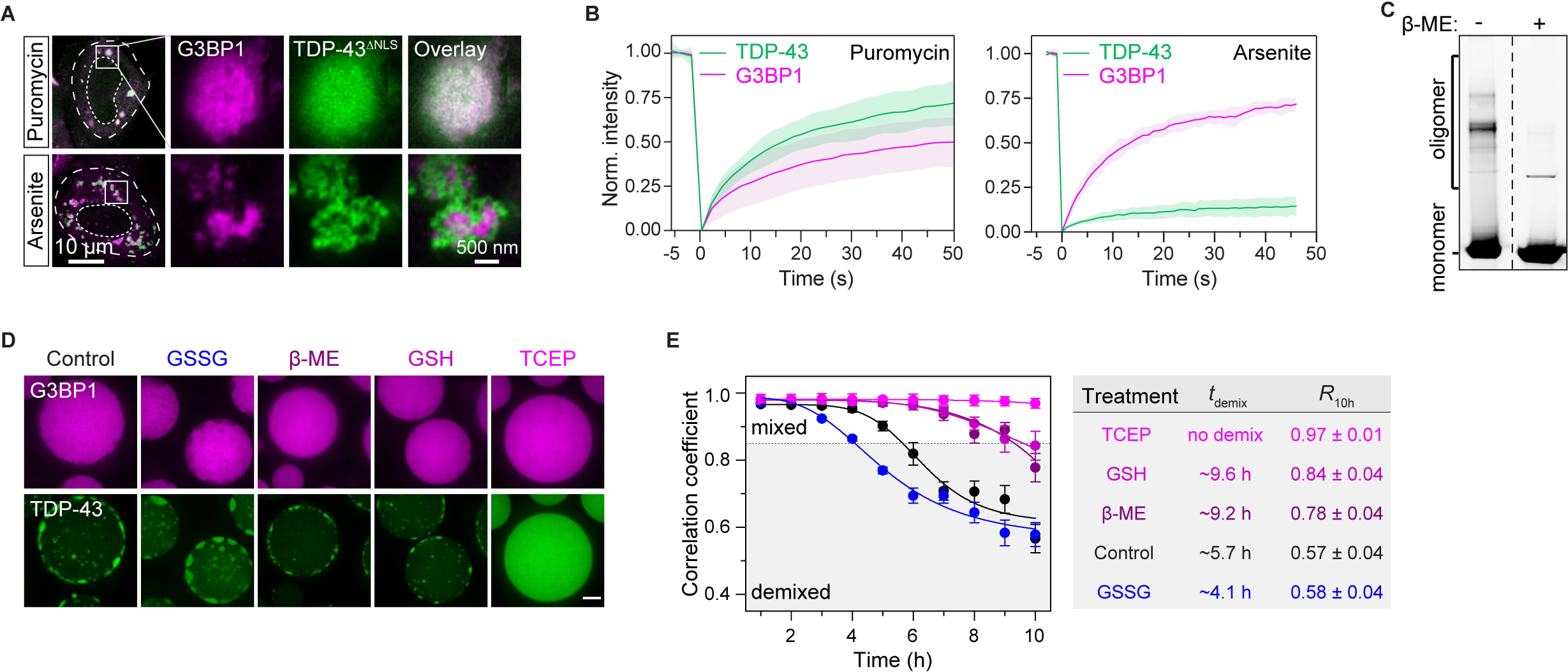
Oxidation is a prerequisite for intra-condensate demixing of TDP-43 (A) Oxidization on intra-condensation demixing of TDP-43 in HeLa cells assayed by STED. Cells expressing TDP-43^ΔNLS^ were treated with 100 μM arsenite or 10 μg/ml puromycin for 120 min. Scale bar, 10 μm and 500 nm for confocal and STED images, respectively. (B) FRAP of TDP-43^ΔNLS^ and G3BP1 in stress granules in HeLa cells after addition of 10 μg/ml puromycin for 180 min or 100 μM arsenite for 60 min. Data represent the mean ± SD. (C) Disulfide bond formation of TDP-43 in minimal stress granules. Stress granules with demixed TDP-43 at 10 h were dissolved with SDS-containing loading buffer and loaded onto SDS-PAGE in the absence or presence of β-ME. (D and E) Redox modifications governing intra-condensate demixing of TDP-43 in minimal stress granules. GSSG (1 mM), β-ME (10 mM), GSH (10 mM) or TCEP (10 mM) were used to treat the stress granules and representative images at 10 h are shown (D). The demixing kinetics, including the demixing time (*t*demix) and correlation coefficient at 10 h (*R*10h), were quantified (E). Data represent the mean ± SD. Scale bar, 10 μm. See also Figure S3.

To directly investigate the role of oxidation on intra-condensate demixing, we turned to the reconstituted minimal stress granules. When subjected to non-reducing SDS-PAGE, demixed TDP-43 in the reconstituted stress granules formed high-molecular-weight oligomers that could be disassembled by adding the reducing agent β-mercaptoethanol (β-ME) (**Figure 3C**), revealing the formation of intermolecular disulfide bonds. When reconstituted stress granules were incubated with the oxidizing agent glutathione disulfide (GSSG), TDP-43 demixing was accelerated and occurred at lower threshold concentrations (**Figures 3D**, **3E** and **S3C**). In contrast, the reducing agents β-ME or glutathione (GSH) slowed down TDP-43 demixing, whereas TCEP, a more potent reducing agent, abrogated demixing (**Figures 3D** and **3E**). Importantly, this suggests that TDP-43 is highly sensitive to redox conditions and that background ROS levels are sufficient to drive TDP-43 demixing in test tube experiments with purified proteins.

Taken together, the data from cells and *in vitro* analysis indicate that TDP-43 aggregation requires a double event: 1) up-concentration of TDP-43 by partitioning into stress granules, and 2) demixing coupled to oxidation-driven misfolding of TDP-43 to promote the assembly of protein aggregates.

### Hydrophobic patch interactions and disulfide bond formation promote homotypic TDP-43 interactions that govern intra-condensate demixing

To generate molecular hypotheses for the mechanisms governing TDP-43 demixing, we turned to molecular dynamics (MD) simulations. TDP-43 is composed of a folded N-terminal domain (NTD), two tandem folded RNA-recognition motifs (RRM1 and RRM2), and a low-complexity C-terminal domain (CTD) (**Figure 4A**). The CTD contains four sub-regions, one being the hydrophobic patch region (HP, 318–343 aa), which adopts a unique α-helical structure^60–62^ (**Figure 4A**).

**Figure 4.**
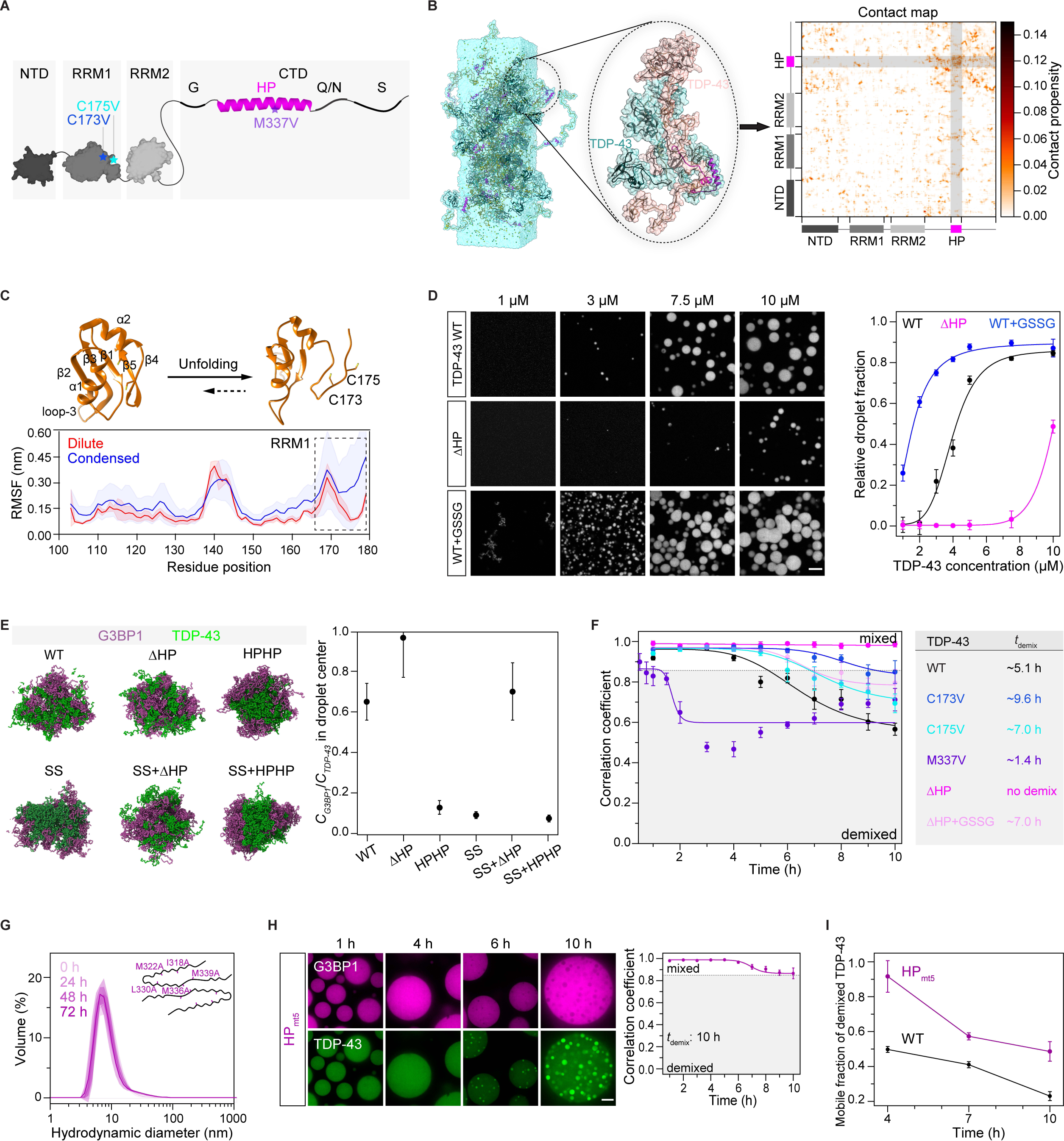
Hydrophobic patch interactions and disulfide bond formation constitute homotypic TDP-43 interactions that govern intra-condensate demixing (A) Schematic of folded and disordered regions in TDP-43 (1–414 aa). NTD, N-terminal domain (1–80 aa); RRM1 and RRM2, RNA recognition motif 1 (106–176 aa) and 2 (191–262 aa); CTD, C-terminal domain (263–414 aa); G, glycine-rich (274–314 aa); HP, hydrophobic patch region (318–343 aa); Q/N, Q and N-rich (344–365 aa); S, serine-rich (370–402 aa). Point mutations studied in the following sections are also shown here. (B) Representative snapshot from an atomistic MD simulation (3 µs) of a full-length TDP-43 condensate composed of 25 chains using a slab-geometry setup (left), close-up of two interacting TDP-43 chains within the condensate (middle) and the corresponding two-dimensional, intermolecular contact map showing pairwise, residue-level interactions (right). In the contact map, contact propensity for a specific residue refers to the average number of contacts (<NIJ>) per frame of the trajectory summated over the pairwise contributions of all TDP-43 chains and normalized by the total number of chains (n=25). (C) Structures of folded (PDB: 4IUF) and partially unfolded RRM1 domain in simulation of the TDP-43 condensed phase (top), and comparison between per-residue root-mean-square fluctuation (RMSF) for RRM1 in the dilute and condensed phases (bottom). The RRM1 domains are shown in ribbon representation with cysteine residues (173/175) shown as sticks. RMSF analysis for each residue, which is used to quantify the local flexibility of proteins, was computed over three independent trajectories for the monomer in the dilute phase (4.5 μs each) and over 25 chains in the condensed phase (2.5 μs each), respectively. Data represent the mean ± SD. (D) The HP interactions and disulfide bond formation favor TDP-43 condensation. Phase separation of TDP-43 WT or ΔHP with titrated concentrations in the absence or presence of GSSG (5 mM) are shown. Relative droplet fraction of TDP-43 versus the total protein concentration is quantified and shown as mean ± SD. Scale bar, 10 μm. (E) Cross-section of TDP-43/G3BP1 droplet snapshots (left) and protein concentration ratio (*CG3BP1*/*CTDP-43*) at the droplet center obtained from the simulations (right). The droplets comprised of an equal number of TDP-43 and G3BP1 chains (25 each). Simulation was computed over 5 blocks of 1–1.1 µs duration after excluding the initial 2 μs as equilibration. A ratio of 1 corresponds to a well-mixed droplet while values approaching 0 indicate strong demixing. Data represent the mean ± SEM. (F) Intra-condensate demixing of TDP-43 variants (10 μM) in minimal stress granules. Pearson colocalization between G3BP1 and TDP-43 was analyzed and the demixing time (*t*demix) is shown. Data represent the mean ± SD. (G) DLS assay for TDP-43 HPmt5. The mutated residues are shown based on the cryo-EM structure of the TDP-43 amyloid cross-β core (PDB: 6n37). HPmt5 (80 μM) was incubated in 500 mM KCl. At the time indicated, samples (5 μM) were assayed by DLS. Data represented as the mean ± SD. (H) Intra-condensate demixing of TDP-43 HPmt5 in minimal stress granules. HPmt5 (10 μM) was added into minimal stress granules. Pearson colocalization between G3BP1 and TDP-43 was analyzed. Data represent the mean ± SD. Scale bar, 10 μm. (I) FRAP assay for mobile fractions of TDP-43 WT and HPmt5 within the demixed phase along intra-condensate demixing in minimal stress granules. Data represent the mean ± SD. See also Figure S4.

Considering intra-condensate demixing generates TDP-43 enriched phase in stress granules, we first carried out all-atom MD simulation on TDP-43 condensates to investigate the underlying homotypic interactions (see STAR Methods). Analysis of pairwise, intermolecular contacts within the condensed phase revealed significant interactions from the disordered CTD, especially the HP region (**Figures 4B** and **S4A**), which adopted an α-helical structure consistent with previous NMR studies^63^ (**Figure S4B**). Importantly, MD simulation also showed that RRM1, but not RRM2, partially unfolds in the β4– β5 region (**Figures 4C, S4C** and **S4D**) in the condensed phase, leading to solvent exposure of cysteine 173 and 175 (**Figure 4C**). The exposure of these cysteine residues would favor disulfide bond formation, and thereby reduce the threshold concentration for condensation *via* homotypic TDP-43 interactions, in line with the oxidation requirement for TDP-43 demixing (**Figures 3A** and **3D**).

To test these simulations, we looked at phase separation of purified TDP-43, in the absence of RNA. Deletion of the HP region significantly decreased TDP-43 condensation, while cysteine oxidation by GSSG increased TDP-43 condensation (**Figure 4D**). Thus, HP interactions in the C-terminal domain and disulfide bond formation in RRM1 collectively represent the main homotypic interactions of TDP-43.

We next investigated the relationship between homotypic interactions of TDP-43 and its intra-condensate demixing. Using coarse-grained MD simulations of TDP-43/G3BP1 co-condensates (see STAR Methods), we found that the strength of homotypic interactions of TDP-43, mediated by HP interactions and disulfide bond formation, positively correlates with TDP-43 demixing from G3BP1 (**Figure 4E**).

To test predictions from the simulations, we generated mutants in which either cysteines at position 173 or 175 in RRM1 were replaced with valine or isoleucine (see STAR Methods). MD simulations and NMR analysis showed that the stability of these variants was comparable to wild-type TDP-43 (**Figures S4E** and **S4F**). Cysteine variants also preserved RNA binding similar to wild-type TDP-43 and exhibited normal RNA processing function in cells as revealed by autoregulation^64^ (**Figures S4G** and **S4H**). In reconstituted stress granules, both the cysteine variants C173V and C175V showed reduced demixing compared to wild-type TDP-43 (**Figures 4F** and **S4I**). The disease variant M337V, previously shown to strengthen the HP interactions,^65^ displayed increased demixing (**Figures 4F** and **S4I**). The ΔHP variant, with significantly reduced demixing in the simulation, could not demix (**Figures 4F** and **S4I**). However, oxidation could trigger the demixing of the ΔHP variant (**Figures 4F** and **S4I**), suggesting a synergy between disulfide bond formation and HP interactions in driving demixing.

We conclude that the interactions mediated by HP interactions and cysteine oxidation in the RRM1 collectively enable TDP-43 phase separation *via* homotypic interactions, thus giving rise to intra-condensate demixing inside stress granules.

### TDP-43 aggregation is driven by an α-helix-to-β-sheet transition of the hydrophobic patch region

Previous work has suggested that irreversible aggregation requires an α-helix to a β-sheet transition of the HP region.^25,26^ To test whether this transition is also required for generating pathological aggregates after intra-condensate demixing, we generated an HP variant (HPmt5), in which five mutations were introduced into the HP to destabilize the amyloid cross-β core (PDB: 6n37).^26^ Compared to wild-type TDP-43, which gradually forms oligomers over time as assayed by dynamic light scattering (DLS) (**Figure S4J**), HPmt5 did not oligomerize (**Figure 4G**). Remarkably, when the HPmt5 protein was left in the test tube for 72 h without oligomerization, it continued to form condensates but not aggregates (**Figure S4K**), suggesting that it retains the α-helical structure critical for condensation. In reconstituted stress granules, HPmt5 could demix (**Figure 4H**), but the liquid-to-solid transition of the demixed phase was significantly delayed compared to wild-type TDP-43 (**Figure 4I**). This allows us to conclude that intra-condensate demixing and the subsequent liquid-to-solid transition are two distinct steps, with helix-to-sheet transition of the HP region required for aggregation in the demixed TDP-43 phase.

### Oxidation-resistant TDP-43 variants with lowered self-assembly propensity abrogate pathological demixing *in vivo*

To leverage the insights gleaned from our *in silico* predictions and *in vitro* studies, we engineered TDP-43^ΔNLS^ variants with altered self-assembly propensities and expressed them in HeLa cells. Supporting the idea that disulfide bond formation and HP interactions synergize to promote TDP-43 demixing, either cysteine mutations or deletion of the HP alone reduced demixing, while the disease mutation M337V enhanced demixing (**Figures 5A** and **S5A**). Importantly, combining the two mutations (denoted as C173V/ΔHP and C175V/ΔHP) abrogated intra-condensate demixing (**Figures 5A** and **5B**). More specifically, although the double mutants were still strongly enriched inside stress granules, they remained dispersed for the time course of the experiment. Consistent with loss of demixing, these variants also did not co-stain with HSPB1 and ubiquitin (**Figures 5C** and **5D**), suggesting that they cannot form pathological aggregates.

**Figure 5.**
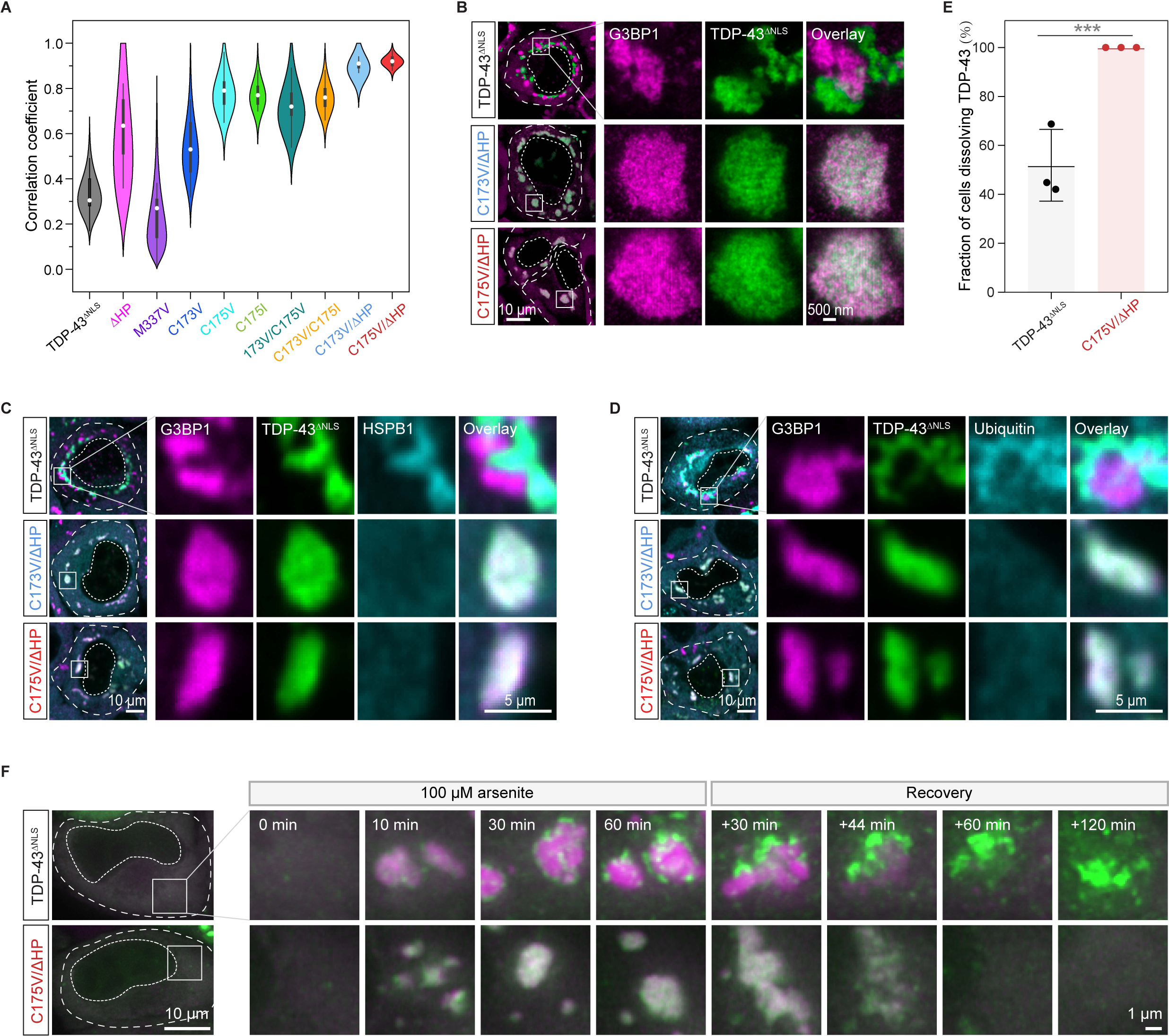
Oxidation-resistant TDP-43 variants with lowered self-assembly propensity abrogate pathological demixing *in vivo* (A) Pearson colocalization between G3BP1 and TDP-43^ΔNLS^ variants under 100 μM arsenite stress for 120 min. (B) Representative STED images of HeLa cells expressing TDP-43^ΔNLS^ variants after addition of 100 μM arsenite for 120 min. Stress granules marked by squares under confocal microscopy were further visualized by STED microscopy. Scale bar, 10 μm and 500 nm for confocal and STED images, respectively. (C and D) Prevention of intra-condensate demixing of TDP-43 abolishing the formation of pathological aggregates. Cells expressing TDP-43^ΔNLS^ variants were stressed with 100 μM arsenite for 120 min and images were acquired for HSPB1 (C) and ubiquitin (D) by confocal microscopy. Stress granules marked by squares were zoomed in. Scale bar, 10 μm and 5 μm for confocal and zoomed in images, respectively. (E) Quantification of dissolution of TDP-43^ΔNLS^ aggregates. HeLa cells expressing TDP-43^ΔNLS^ variants were stressed with 100 μM arsenite for 60 min following 120 min recovery. The demixed TDP-43^ΔNLS^ aggregates were monitored in cells capable of dissolving stress granules, and data were presented as the fraction of these cells able to dissolve TDP-43^ΔNLS^ aggregates. Data represent the mean ± SD. (F) Live imaging of cells expressing TDP-43^ΔNLS^ variants during stress with 100 μM arsenite for 60 min and recovery for 120 min. Stress granules marked by squares were zoomed in. Scale bar, 10 μm and 1 μm for fluorescent and zoomed in images, respectively. See also Figure S5, Videos S5 and S6.

ALS is thought to be caused in large parts by a gain of toxicity from persisting protein aggregates.^17,66^ To investigate whether intra-condensate demixing leads to aggregates that persist, we followed their stability after stress removal. Of those cells in which G3BP1 condensates dissolved, TDP43^ΔNLS^ aggregates remained in about 50% of cells (**Figures 5E**, **5F**, and **Video S5**). Importantly, in cells expressing C175V/ΔHP, where intra-condensate demixing was not observed, all cells were able to dissolve TDP-43 assemblies (**Figures 5E**, **5F**, and **Video S6**). We were able to use these assays to distinguish the distinct roles of cysteines and the HP region in forming irreversible aggregates. Variants in which the HP region was deleted (ΔHP) or mutated (HPmt5) significantly reduced recruitment of aggregation markers such as HSPB1 and ubiquitin (**Figures S5B** and **S5C**). After stress removal, they were also dissolved better compared to wild-type and particularly the disease variant M337V (**Figure S5D**). The combined data suggest that although demixing can be driven by disulfide bond formation, stable aggregation requires the helix-to-sheet transition of HP region within demixed condensates. The data further illustrate how pathological aggregates can emerge from dynamic multicomponent condensates and persist even after removal of stress. Such aggregates could be toxic and require specific and energy-intensive mechanisms for their removal.^33^

### Intra-condensate demixing results in pathological TDP-43 aggregates in motor neurons, accompanied by disruption of nucleocytoplasmic transport

To assess the pathological relevance of intra-condensate demixing, we expressed TDP-43^ΔNLS^ in iPSC-derived motor neurons (iPS-MN), a more physiological cell line affected by TDP-43 pathology. Upon arsenite stress, TDP-43^ΔNLS^ was initially enriched and well dispersed inside stress granules, but later demixed into small puncta (**Figure 6A** and **Video S7**), thus highlighting intra-condensate demixing as a universal phenomenon across different cell lines. In contrast, puromycin was unable to trigger TDP-43 demixing (**Figure 6B**), emphasizing a similar oxidation requirement in motor neurons. Accompanying the demixing process induced by arsenite, TDP-43 dynamics gradually slowed down by FRAP, while TDP-43 stayed dynamic under puromycin stress (**Figure 6C**), indicating hardening during intra-condensate demixing. These demixed puncta co-stained with HSPB1 and exhibited ubiquitination as well (**Figure 6D**). Therefore, intra-condensate demixing generates TDP-43 aggregation with pathological hallmarks in motor neurons.

**Figure 6.**
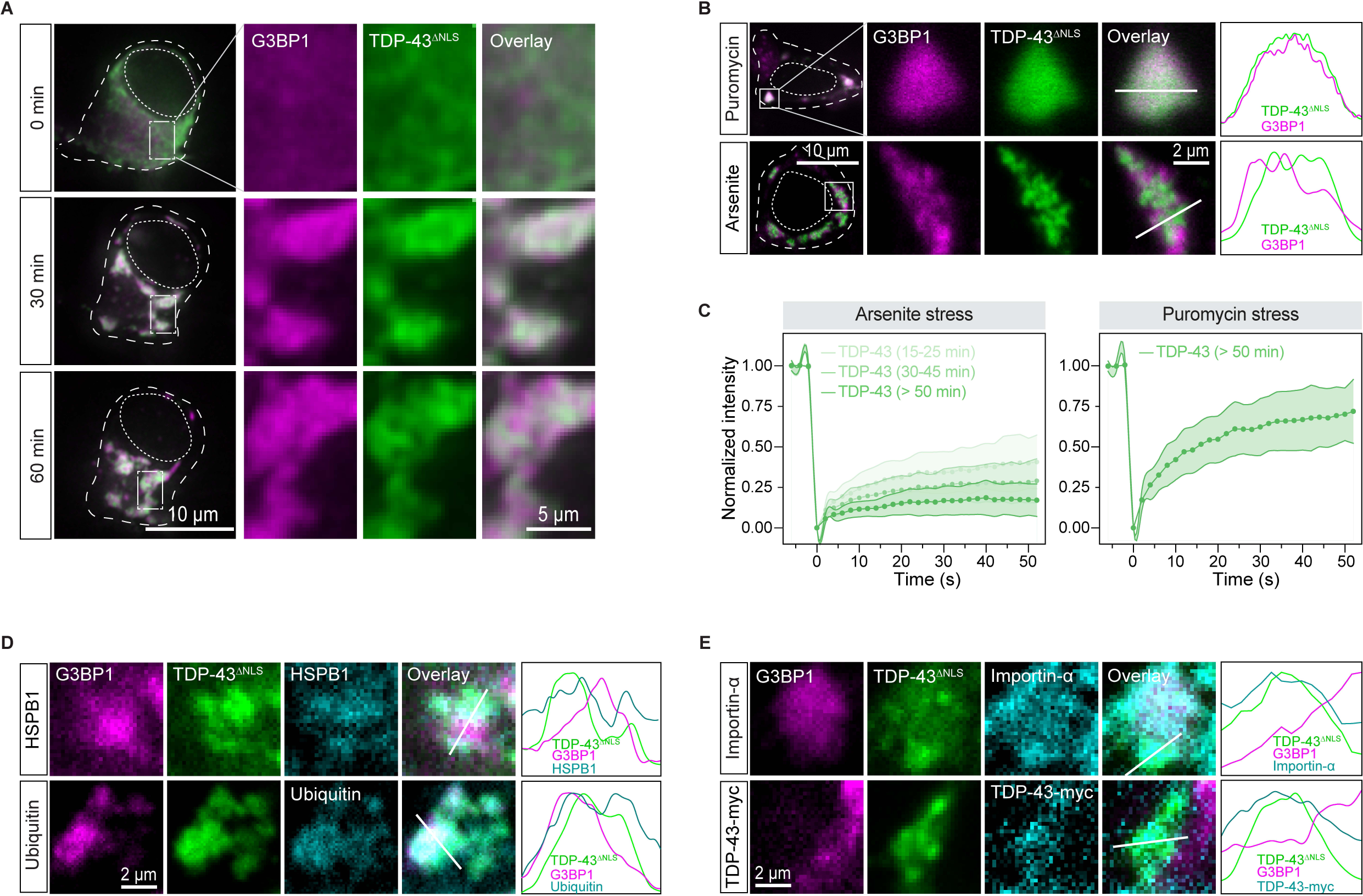
Intra-condensate demixing results in pathological aggregates in motor neurons, accompanied by impairment of nucleocytoplasmic transport (A) Representative images of intra-condensate demixing of TDP-43^ΔNLS^ in iPS-MN. The cells were cotransfected with GFP-tagged TDP-43^ΔNLS^ and mCherry-tagged G3BP1 and live-cell imaging was performed after addition of 100 μM arsenite. Scale bar, 10 μm and 5 μm for whole-cell and zoomed in images, respectively. (B) Oxidization on intra-condensation demixing of TDP-43 in iPS-MN cells. Motor neurons expressing TDP-43^ΔNLS^ were treated with 10 μg/ml puromycin or 100 μM arsenite for 60 min. Normalized fluorescent intensities for TDP-43 and stress granules along the straight line are shown. Scale bar, 10 μm and 2 μm for confocal and zoomed in images, respectively. (C) FRAP of TDP-43^ΔNLS^ in stress granules in iPS-MN cells after addition of 100 μM arsenite or 10 μg/ml puromycin for 60 min. Data represent the mean ± SD. (D) TDP-43 demixing generating pathological aggregates. Motor neurons expressing TDP-43^ΔNLS^ were stressed with 100 μM arsenite for 90 min, and images were acquired for HSPB1 or ubiquitin staining by confocal microscopy. Normalized fluorescent intensities for each channel along the straight line are shown. Scale bar, 2 μm. (E) TDP-43 aggregation impairing nucleocytoplasmic transport in iPS-MN. Motor neurons expressing TDP-43^ΔNLS^ alone or TDP-43^ΔNLS^ together with TDP-43-myc with intact NLS were stressed with 100 μM arsenite for 90 min, and images were acquired for importin-α or TDP-43-myc staining by confocal microscopy. Normalized fluorescent intensities for each channel along the straight line are shown. Scale bar, 2 μm. See also Video S7.

Impairments in nucleocytoplasmic transport have been associated with pathology in ALS patients.^67,68^ We next investigated whether TDP-43 aggregation generated in stress granules causes pathology similar to that observed in patients. TDP-43^ΔNLS^ aggregates were observed to exhibit interactions with the essential nuclear transport factor importin 2α (**Figure 6E**). Supporting the disturbed nucleocytoplasmic transport, demixed TDP-43^ΔNLS^ sequestered nuclear wild-type TDP-43 with the intact NLS. Therefore, TDP-43 aggregates in condensates may render toxicity by sequestering key cellular factors.

## DISCUSSION

In this work, we have investigated the role of stress granules in generating cytosolic TDP-43 aggregates, which are the most prevalent pathological hallmark of ALS and FTD. The key result of our study is that TDP-43 aggregation in cells requires a double event. One event is the increase in TDP-43 concentration within stress granules that surpasses a critical threshold, and the other is oxidative stress, which increases the tendency of TDP-43 to self-interact. These two events collectively trigger intra-condensate demixing of TDP-43 inside stress granules, and this highly concentrated TDP-43 phase undergoes a liquid-to-solid transition. By uncovering the molecular pathway underlying TDP-43 demixing, we were able to abrogate TDP-43 aggregation in cells. Targeting intra-condensate demixing of TDP-43 within stress granules, as a distinct step on the pathway to aggregation, thus becomes a potential avenue for therapeutic intervention.

Combining our data with previous results allows us to define the role of stress granules in TDP-43 aggregation, which is summarized in Figure 7. Under physiological conditions, TDP-43 is enriched in the nucleus and bound to its RNA targets^16,69^ (step 1). However, during oxidative stress, TDP-43 passively leaks into the cytoplasm, where it partitions into stress granules^37^ (step 2). Stress granules recruit TDP-43 by providing RNA binding sites, promoting its gradual up-concentration (**Figure 1A**). Once TDP-43 crosses a critical threshold concentration (*Cthreshold*) under oxidation conditions, the stress granule condensate formed by a network of heterotypic interactions undergoes intra-condensate demixing (step 3a), giving rise to a separate but initially dynamic TDP-43 phase (step 3b) (**Figures 1B**, **2B** and **6A**). This process is primarily driven by homotypic interactions among TDP-43 molecules mediated by transient, α-helical HP interactions and disulfide bond formation in the RRM1 (step 3c), which disfavors heterotypic interactions of TDP-43 with RNA and promotes homotypic TDP-43 interactions (**Figures 4B, 4C and 4F**). The demixed phase facilitates a liquid-to-solid transition into aggregates exhibiting pathological hallmarks of TDP-43 inclusions (step 4a) (**Figures 1E**, **1F** and **6D**). The aggregation is likely mediated by stable cross-β-sheet interactions from the HP region (step 4b) (**Figures 4H** and **4I**). Further structural work will be required to study the mechanism of aggregation.

**Figure 7.**
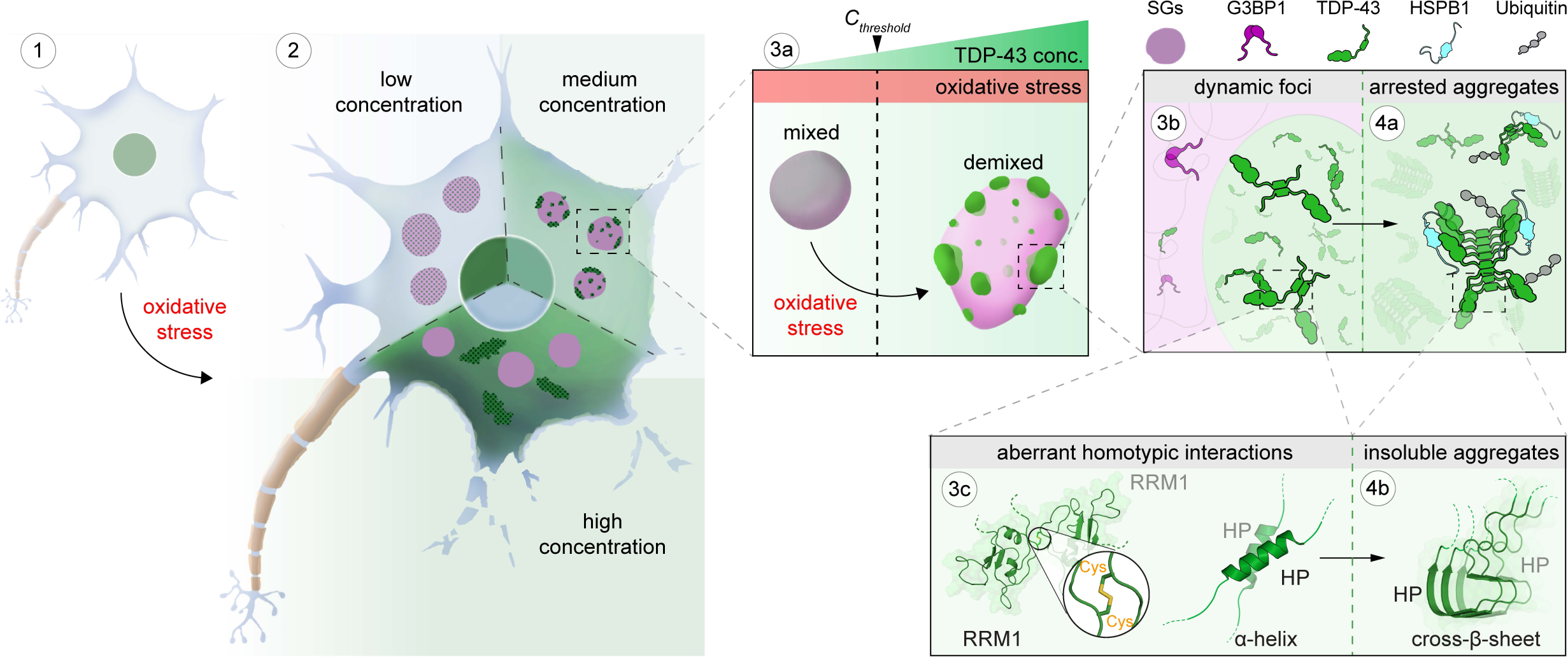
Model of intra-condensate demixing of TDP-43 inside stress granules generating pathological aggregates The intra-condensate demixing of TDP-43 inside stress granules generating pathological aggregates at different scales, from cells to condensates to molecular basis, is shown schematically. Intra-condensate demixing of TDP-43 occurs by up-concentration inside stress granules under oxidation (step 3a). The demixed but dynamic TDP-43 phase facilitates a liquid-to-solid transition into pathological aggregates (steps 3b and 4a). See Discussion for details.

Based on cell culture, biochemical assays, MD simulations as well as previously published data, we can build a molecular picture of how homotypic interactions of TDP-43 are favored in the heterotypic environment of stress granules. Raising the local concentration of TDP-43 above a critical threshold by partitioning into stress granules is a prerequisite for the homotypic interactions that drive TDP-43 demixing (**Figures 1A** and **2C**), presumably because it increases the proximity of TDP-43 molecules.^70^ Simulations suggest that the interactions are potentiated by two molecular events: One event is partial unfolding of RRM1, which exposes cysteine residues in RRM1 (**Figure 4C**). Oxidative stress, shown to play crucial roles in the progression of various neurodegenerative diseases,^71^ increases disulfide bond formation of exposed cysteine residues (**Figures 3A**, **3D** and **6B**). The other event is α-helical interactions between the C-terminal HP region^63^ (**Figure 4B**). Indeed, disease mutations within the HP region that favor homotypic interactions can promote TDP-43 demixing (**Figures 4F** and **5A**). The requirement for oxidation and the exacerbation by disease mutations suggest that the intra-condensate demixing of TDP-43 is relevant in the pathogenesis of neurodegenerative diseases.^29,53^

Our data suggest why it has been difficult to pinpoint the trigger for TDP-43 aggregation in cells. Presumably, TDP-43 also misfolds and forms disulfide bonds in the surrounding cytosol under oxidative stress, but the probability for misfolded TDP-43 molecules is not sufficient to trigger homotypic interactions at physiological concentrations. By locally raising the concentration, intra-condensate demixing within stress granules is required to promote the assembly of persistent pathological TDP-43 aggregates. This is likely because the highly concentrated demixed phase can increase the probability of a helix-to-sheet conversion of the HP region (**Figures 4G–4I**), which eventually facilitate the transition of low-complexity CTD into cross-β-sheet structures.^25,72^ These TDP-43 aggregates generated inside stress granules appear to be resistant to removal by the protein quality control machinery, as demonstrated by experiments looking at recovery from stress (**Figure 5F**). We speculate that this may be due to tight cross-β interactions and the formation of an extensive network of disulfide bonds, but this requires additional work in the future.

Multiphasic condensate architectures have been widely described in a cellular context, including nucleoli, nuclear speckles, paraspeckles and P granules.^73–76^ The demixed state in multiphasic condensates is fundamentally underpinned by differences in the molecular grammar of individual scaffold molecules governing each phase.^74,77^ Presumably these multiphase architectures are additionally dependent upon specific interactions between their diverse set of constituents.^35,78^ Here we propose that, in the context of disease, initially well-mixed, multicomponent condensates such as stress granules can similarly undergo a transition into multiphasic condensates, giving rise to a distinct homotypic phase inside a complex heterotypic system that promotes disease emergence.

## Supporting information

Video S1 Low expression of TDP-43 dNLS dispersed inside stress granules in HeLa cells

Video S2 High expression of TDP-43 dNLS forming aggregates independent of stress granules in HeLa cells

Video S3 Medium expression of TDP-43 dNLS undergoing intra-condensate demixing inside stress granules in HeLa cells

Video S4 Intra-condensate demixing of TDP-43 inside minimal stress granules

Video S5 Dissolution assay of demixed TDP-43 dNLS aggregates in HeLa cells

Video S6 Dissolution assay of TDP-43 dNLS C175V dHP inside stress granules in HeLa cells

## ACKNOWLEDGMENTS

We would like to acknowledge technical support by E. Geertsma, B. Borgonovo, R. Lemaitre and A. Bogdanova (PEPC facility); A. Pozniakovski and L.D. Mamede (DNA constructs); J. Peychl, and B. Schroth-Diez (light microscopy facility); R. Barsacchi, M. Stöter and C. Möbius (technology development studio); C. Eugster Oegema (organoid and stem cell facility). We would like to thank P. McCall and T. Harmon for discussions on theory. We would like to thank Y. Ye for the help on the cartoon model. We would like to thank A. Karasavvidi for her contributions to confocal microscopy experiments and E.H. Hemamali for performing the RNA binding assays. We would like to thank J. Wang, C. Hoege, A. Fritsch, S. Maharana, J Guillén-Boixet, H. Uechi, A. Majumdar, A. Klosin, L. Hubatsch, A. Pal, M. Ruer-Gruß, J. Savage and D. Sun for critical, as well as very fruitful discussions. We acknowledge funding from the Max Planck Society and the European Research Council (PhaseAge, ERC grant agreement number 725836) to S.A. and A.A.H. Funding from Korber Stiftung to A.A.H. Funding by Volkswagen Foundation to D.K. and A.A.H. National Institutes of Health (NIH) National Institute for Neurological Disorders and Stroke (NINDS) and National Institute on Aging (NIA) R01 NS114289, and the Department of Defense CDMRP/ALSRP W81XWH-20-1-0241 to Y.M.A.

## AUTHOR CONTRIBUTIONS

A.A.H., S.A. and X.Y. designed and coordinated the project. X.Y. planned and co-performed all of the experiments. D.K. planned and performed biochemical experiments and reconstitution assays with initial observation of intra-condensate demixing *in vitro* and its oxidation-dependence. J.N. performed time-resolved confocal microscopy experiments with initial observations of dynamicity, concentration-dependence, and dissolution phenotypes in *vivo*. K.P.-G. performed stimulated emission depletion microscopy with initial observation of intra-condensate demixing *in vivo*. P.M., A.R. and J.M. designed and performed the all-atom MD simulation and Coarse-grained MD simulations. T.M.F and A.S. performed additional biochemical experiments. P.M.P, L.G. and Y.M.A performed RNA binding and TDP-43 autoregulation assay. S.H.W., J.S., H.L.D. and N.L.F. performed NMR experiments to characterize the structure of cysteine variants. A.H. established and provided the STED set-up. A.A.H., S.A., X.Y. and D.K. drafted the manuscript with contributions from other authors. All authors contributed to data analysis and interpretation.

## SUPPLEMENTAL FIGURE LEGENDS

**Figure S1.**
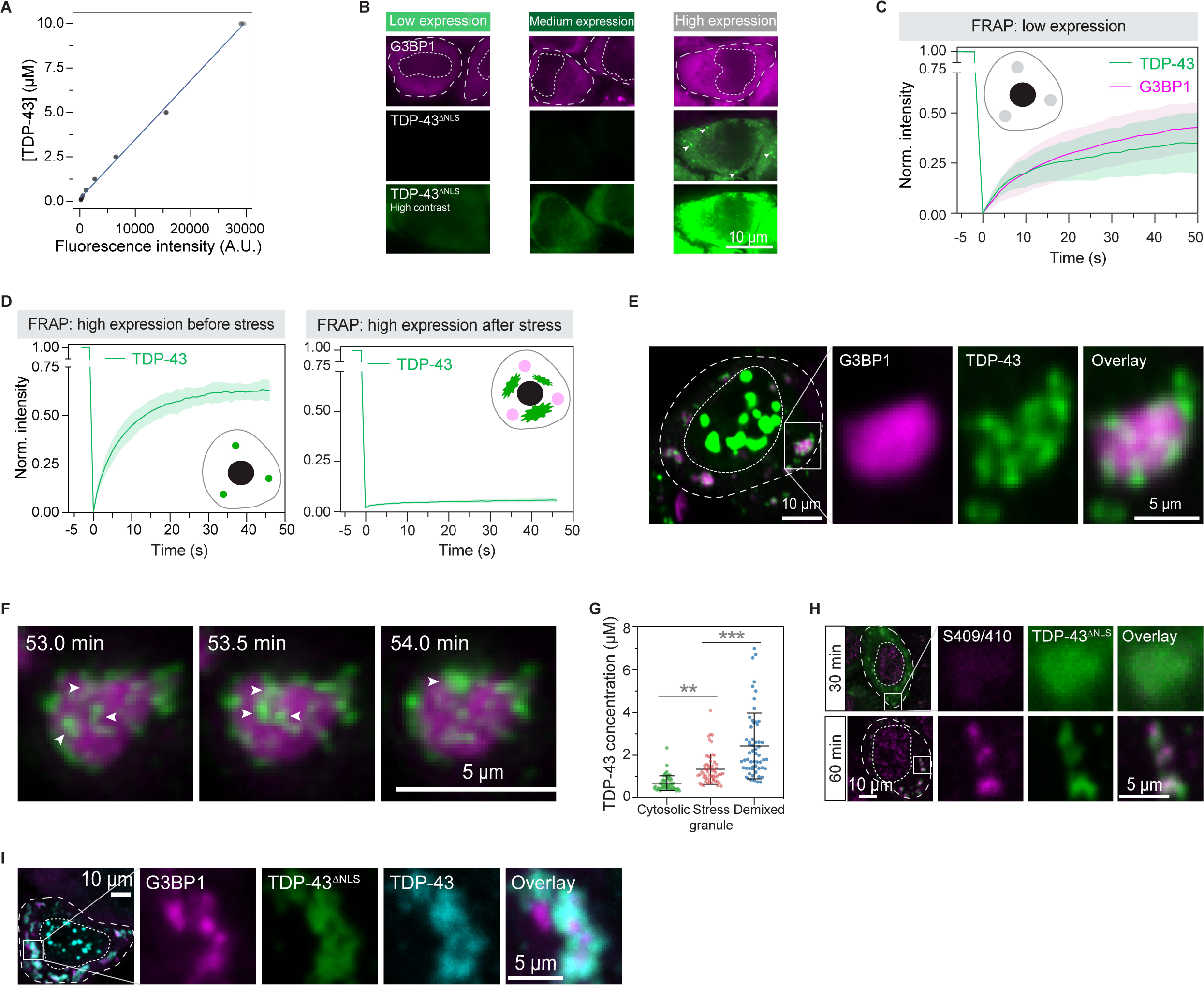
Intra-condensate demixing and aggregate formation of TDP-43 *in vivo*, related to Figure 1 (A) Fluorescent intensity curve of recombinant GFP-tagged TDP-43 at different concentrations. (B) Representative images of HeLa cells with low, medium and high expression of TDP-43^ΔNLS^ before stress during live-cell imaging. High-contrast images are also shown to highlight the low and medium expression cells. The cellular and nucleus boundaries are indicated by dashed lines. Arrow heads indicate small puncta prior to stress in high expression cells. Scale bar, 10 μm. (C) FRAP of TDP-43^ΔNLS^ and G3BP1 in stress granules in low expression cells after addition of 100 μM arsenite for 60 min. Data represent the mean ± SD. (D) FRAP of TDP-43^ΔNLS^ puncta (G3BP1-negative) in high expression cells before stress and after addition of 100 μM arsenite for 60 min. Data represent the mean ± SD. (E) Representative confocal images of HeLa cells expressing wild-type TDP-43 with intact NLS after addition of 100 μM arsenite for 120 min. Scale bar, 10 μm and 5 μm for confocal and zoomed in images, respectively. (F) Representative images of demixed TDP-43^ΔNLS^ puncta fusion during live-cell imaging. Puncta undergoing fusion are indicated by white arrowheads. Scale bars, 5 μm. (G) TDP-43^ΔNLS^ concentrations in HeLa cells in the cytosol before stress (cytosolic), in stress granules at the beginning of demixing (stress granule) and in demixed TDP-43^ΔNLS^ puncta at the end of demixing (demixed). (H) Phosphorylation of TDP-43^ΔNLS^ aggregation upon intra-condensate demixing by confocal microscopy. Cells expressing TDP-43^ΔNLS^ were stressed with 100 μM arsenite and images were acquired before and after demixing. Scale bar, 10 μm and 5 μm for confocal and zoomed in images, respectively. (I) Cytoplasmic TDP-43^ΔNLS^ aggregates generated from intra-condenste demixing recruiting nuclear TDP-43 by confocal microscopy. Cells cotransfected with GFP-tagged TDP-43^ΔNLS^ (100 ng) and Myc-tagged TDP-43 (50 ng) with intact NLS were stressed with 100 μM arsenite for 120 min. Scale bar, 10 μm and 5 μm for confocal and zoomed in images, respectively.

**Figure S2.**
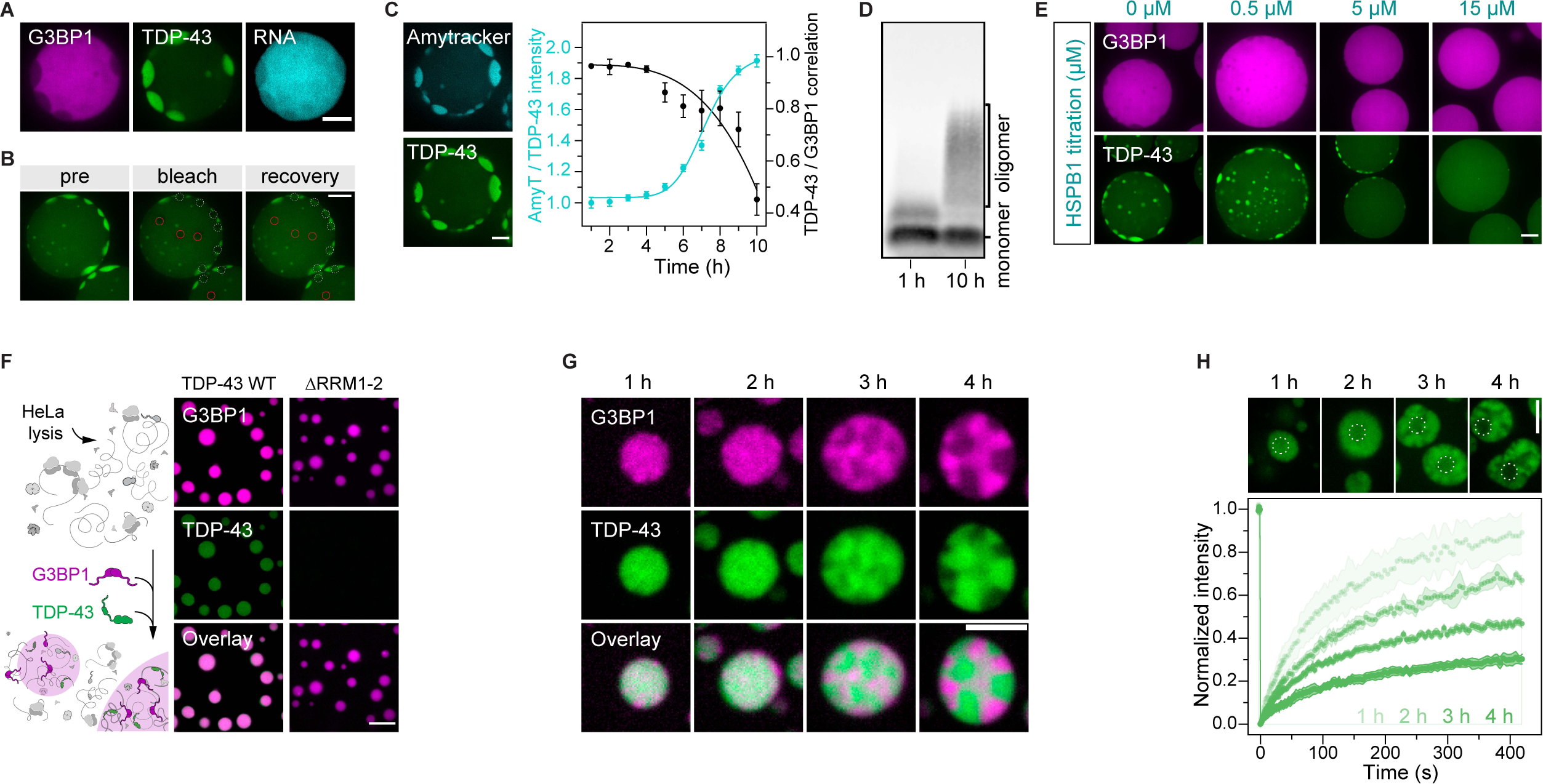
Intra-condensate demixing and aggregate formation of TDP-43 in reconstituted stress granules, related to Figure 2 (A) Reduced RNA binding by TDP-43 upon intra-condensate demixing in minimal stress granules at 10 h. The TDP-43-rich phase contains less RNA stained by BoBo-3 dye (1 μM) than the G3BP1-rich phase. Scale bar for Figure S2, 10 μm. (B) Representative condensates in minimal stress granules at 10 h are shown before and after bleaching. The sites of bleaching for mixed and demixed phases are indicated by red solid and white dashed circles, respectively. (C) The increasing amytracker staining of demixed TDP-43 along intra-condensate demixing process. Amyloid-specific dye amytracker (2.5 μM) was added into minimal stress granules and the ratio of fluorescent intensity between amytracker and TDP-43 was measured to show the increasing stoichiometry. Data represent the mean ± SD. (D) TDP-43 aggregation in minimal stress granules by SDD-AGE assay. Stress granules containing TDP-43 (10 μM) at 1 h and 10 h were dissolved by 0.3% sarkosyl and analyzed by SDD-AGE containing 0.3% sarkosyl. TDP-43 monomer and oligomer are indicated accordingly. (E) Prevention of TDP-43 demixing by HSPB1 in minimal stress granules. The experiment was carried out with titrated concentrations of HSPB1 and representative images at 10 h are shown. (F) TDP-43 recruitment into reconstituted lysate stress granules. Recombinant G3BP1 (20 μM) and lysate from HeLa cells containing ∼2 μg/μl cellular proteins were incubated to form reconstituted lysate stress granules. TDP-43 WT or ΔRRM1-2 (0.5 μM) was included as a client of stress granules. (G) Intra-condensate demixing of TDP-43 in lysate stress granules. TDP-43 (10 μM) was added into lysate stress granules in the presence of 2.5% dextran. (H) FRAP of TDP-43 in reconstituted lysate stress granules as in (G) over time. The bleaching sites upon recovery are indicated by white dashed circles. Data represent the mean ± SD.

**Figure S3.**
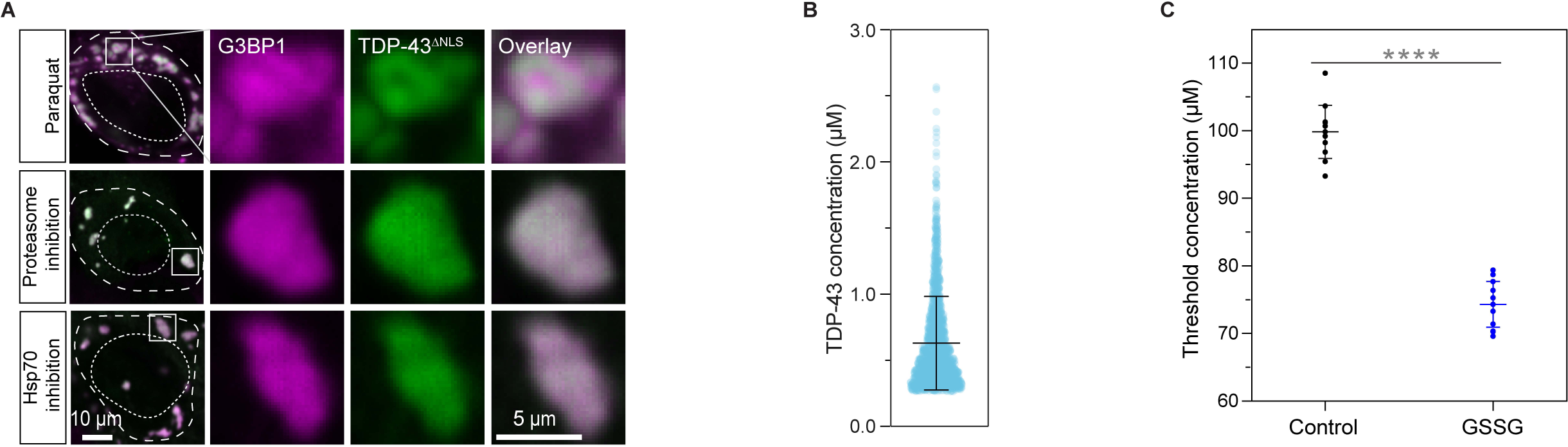
Oxidation is required for intra-condensate demixing of TDP-43, related to Figure 3 (A) Effects of different stressors on intra-condensation demixing of TDP-43^ΔNLS^ in HeLa cells by confocal microscopy. Cells expressing TDP-43^ΔNLS^ were treated with paraquat alone (5 mM) for oxidation, VER-155008 (10 μM) for Hsp70 inhibition or MG132 (10 μM) for proteasome inhibition in the presence of puromycin (10 μg/ml) for 120 min. Scale bar, 10 μm and 5 μm for confocal and zoomed in images, respectively. (B) TDP-43^ΔNLS^ concentrations inside stress granules after addition of puromycin (10 μg/ml) in HeLa cells for 180 min. (C) The threshold concentration for intra-condensate demixing of TDP-43 in minimal stress granules in the absence or presence of GSSG (1 mM).

**Figure S4.**
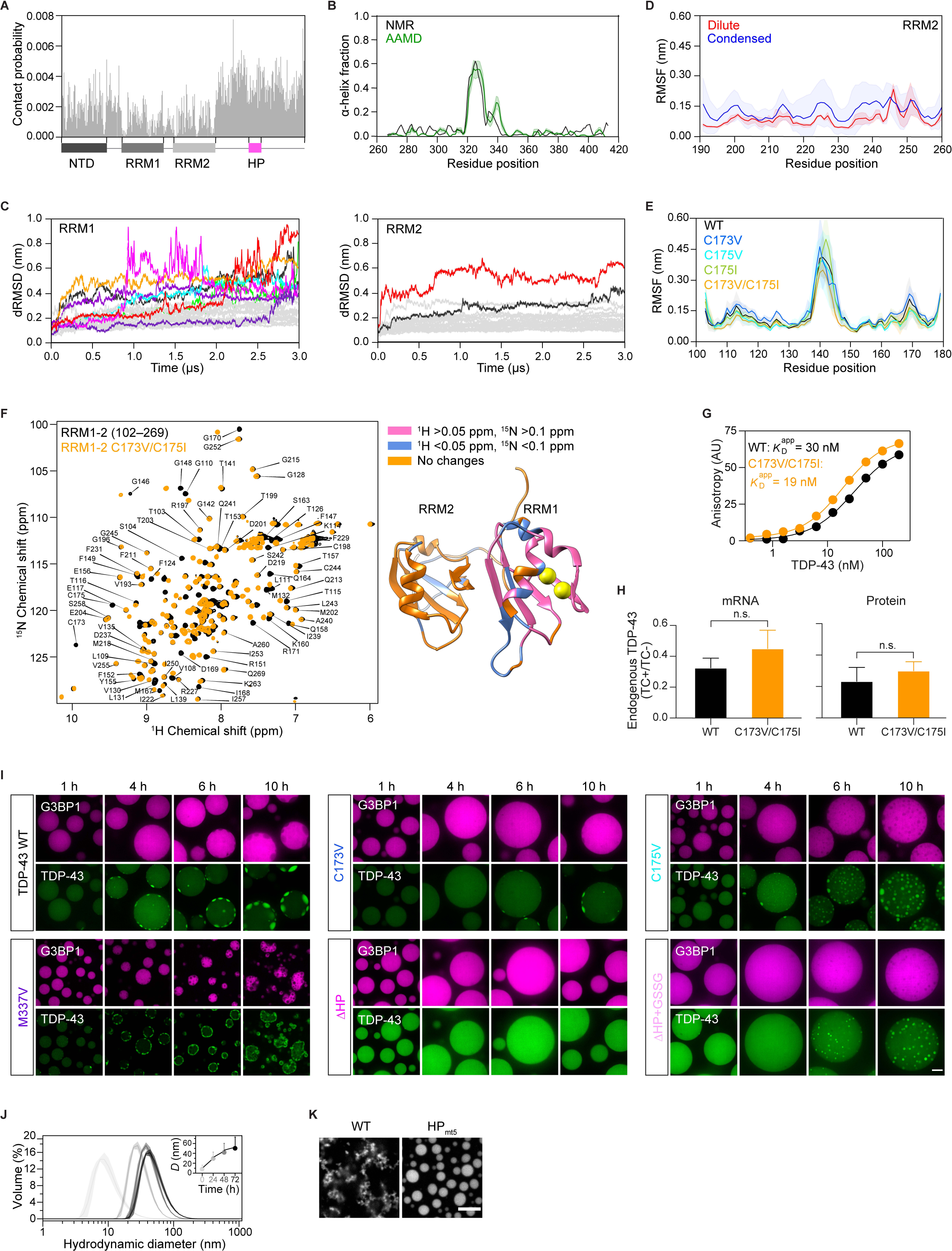
Homotypic interactions mediated by HP interactions and disulfide bond formation govern TDP-43 demixing, related to Figure 4 (A) One-dimensional contact map showing per-residue interaction probabilities which were computed through the summation of all pairwise interactions for each residue position from the two-dimensional contact map in Figure 4B. (B) Comparison of per-residue α-helix fraction for CTD of TDP-43 (267–414 aa) between MD simulations in condensed phase and solution NMR. α-helix fractions were computed based on DSSP secondary structure definitions for MD simulations over 25 chains for the 2.5 µs trajectory, and secondary chemical shifts using the delta-2D program for NMR. Data represent the mean ± SEM. (C) dRMSD of Cα atoms as a function of time with respect to the initial conformation (t=0) for RRM1 (left) and RRM2 (right). Stable RRM domains (dRMSD<0.4 nm) are shown in light gray while unstable domains (dRMSD>0.4 nm) are colored differently. (D) Comparison between per-residue RMSF for RRM2 domain in the dilute and condensed phase from atomistic MD simulations. The mean RMSF was computed over three independent trajectories (4.5 μs each) for the dilute phase (monomer) and 25 chains (2.5 μs each) for the condensed phase, respectively. Data represent the mean ± SD. (E) Conformational stability of RRM1 cysteine variants in comparison to the wild-type based on RMSF analysis at 300 K. RMSF for each residue was calculated over three independent trajectories (2.0 μs each, excluding the first 500 ns). Data represent the mean ± SD. (F) ^1^H-^15^N heteronuclear single quantum coherence (HSQC) spectra of wild-type RRM1-2 (102–269 aa, black) and RRM1-2 C173V/C175I (orange) (left). The spectra were recorded at 298 K using 500 μM protein in 50 mM KPi buffer (pH 6.8) and 150 mM NaCl. Chemical shift assignments were transferred from the BMRB deposited data (BMRB ID: 27613). The chemical shift perturbations of C173V/C175I are mapped on wild-type RRM1-2 (PDB: 4BS2), and 173/175 residues are highlighted in spheres (right). Colors are assigned according to ^1^H and ^15^N changes in residues, with ^1^H >0.05 ppm and ^15^N >0.1 ppm in magenta, ^1^H <0.05 ppm and ^15^N <0.1 ppm in light blue, and residues without substantial change in orange, respectively. (G) Affinity of TDP-43 variants for A(GU)6 RNA, determined by fluorescence anisotropy. FITC-labeled RNA (5 nM) was added into series diluted TDP-43 in Tris buffer (pH 8.0) containing 150 mM NaCl. (H) Autoregulation function of TDP-43 in cells. HEK293 cell lines stably expressing HA-tagged wild-type TDP-43 or C173V/C175I upon tetracycline induction for 72 h. Ratio of endogenous TARDBP mRNA and TDP-43 protein between the presence (TC+) and the absence of induction (TC-) are shown, respectively. Data represent the mean ± SD. (I) Intra-condensate demixing assay for TDP-43 variants. TDP-43 variants (10 μM) were added into the minimal stress granules formed by recombinant G3BP1 (20 μM) and Poly(A) RNA (80 ng/μl) in the presence of 2.5% dextran. Scale bar, 10 μm. (J) TDP-43 oligomer formation by DLS assay. Wild-type TDP-43 (80 μM) was incubated in 500 mM KCl at 25°C. At the time indicated, aliquots of the samples were diluted into 5 μM in 500 mM KCl and assayed by DLS. Data represent the mean ± SD. (K) TDP-43 HPmt5 maintaining droplet formation. Wild-type TDP-43 or HPmt5 (80 μM) was incubated in 500 mM KCl. After 72 h, samples were diluted into a final concentration of 10 μM in 75 mM KCl. Scale bar, 10 μm.

**Figure S5.**
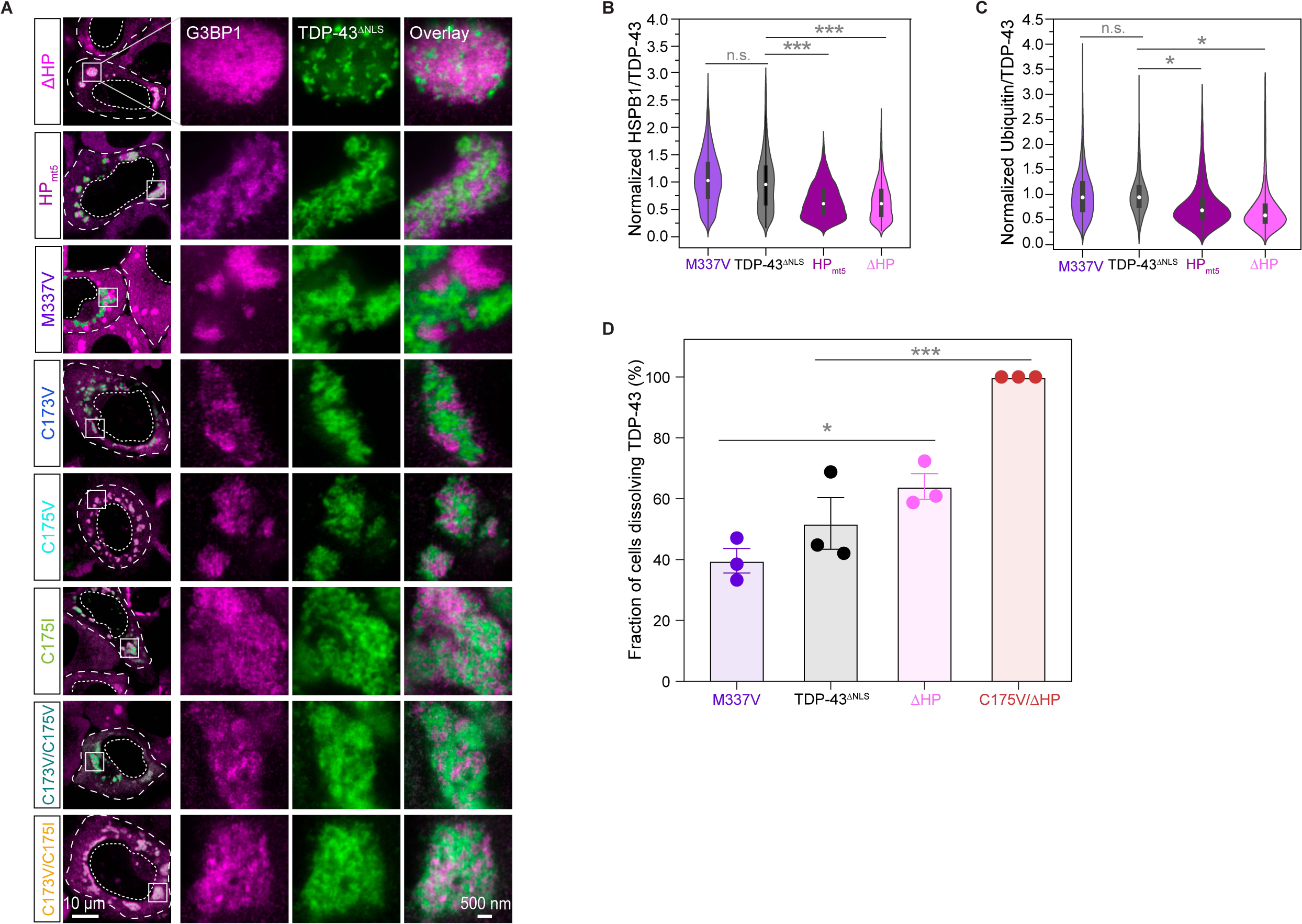
TDP-43 variants with oxidation-resistance and lowered self-assembly propensity abrogate demixing and aggregation *in vivo*, related to Figure 5 (A) Intra-condensate demixing of TDP-43 variants in HeLa cells by STED. Cells expressing TDP-43^ΔNLS^ variants were stressed with 100 μM arsenite for 120 min and images were taken by STED microscopy. The cellular and nucleus boundaries are indicated by dashed lines. Scale bar, 10 μm and 500 nm for confocal and STED images, respectively. (B and C) The HP region interactions contributing to TDP-43 aggregation in the demixed phase. Cells expressing TDP-43^ΔNLS^ variants were stressed with 100 μM arsenite for 60 min, and the ratio of fluorescent intensity between HSPB1 (B) or ubiquitin (C) and demixed TDP-43^ΔNLS^ was quantified. (D) Quantification of dissolution of TDP-43^ΔNLS^ aggregates. HeLa cells expressing TDP-43^ΔNLS^ variants were stressed with 100 μM arsenite for 60 min following 120 min recovery. The demixed TDP-43^ΔNLS^ aggregates were monitored in cells capable of dissolving stress granules, and data were presented as the fraction of these cells able to dissolve TDP-43^ΔNLS^ aggregates. Data represent the mean ± SD.

## SUPPLEMENTAL VIDEO LEGENDS

Video S1. Low expression of TDP-43^ΔNLS^ dispersed inside stress granules in HeLa cells, related to Figure 1

HeLa cells with low expression of TDP-43^ΔNLS^ were stressed by addition of 100 μM arsenite. TDP-43^ΔNLS^ (green) and stress granules marked by mCherry-tagged G3BP1 (magenta) were visualized. Scale bar, 10 μm.

Video S2. High expression of TDP-43^ΔNLS^ forming aggregates independent of stress granules in HeLa cells, related to Figure 1

HeLa cells with high expression of TDP-43^ΔNLS^ were stressed by addition of 100 μM arsenite. TDP-43^ΔNLS^ (green) and stress granules marked by mCherry-tagged G3BP1 (magenta) were visualized. Scale bar, 10 μm.

Video S3. Medium expression of TDP-43^ΔNLS^ undergoing intra-condensate demixing inside stress granules in HeLa cells, related to Figure 1

HeLa cells with medium expression of TDP-43^ΔNLS^ were stressed by addition of 100 μM arsenite. TDP-43^ΔNLS^ (green) and stress granules marked by mCherry-tagged G3BP1 (magenta) were visualized. Scale bar, 10 μm.

Video S4. Intra-condensate demixing of TDP-43 inside minimal stress granules, related to Figure 2

TDP-43 (10 μM) was added into minimal stress granules formed by G3BP1 (20 μM) and Poly(A) RNA (80 ng/μl) in the presence of 2.5% dextran. TDP-43 (green) and G3BP1 (magenta) were visualized. Scale bar, 10 μm.

Video S5. Dissolution assay of demixed TDP-43^ΔNLS^ aggregates in HeLa cells, related to Figure 5

HeLa BAC-G3BP1-mCherry cells with medium expression of TDP-43^ΔNLS^ were stressed by addition of 100 μM arsenite for 60 min and recovery for 120 min. TDP-43^ΔNLS^ (green) and stress granules marked by mCherry-tagged G3BP1 (magenta) were visualized. Scale bar, 10 μm.

Video S6. Dissolution assay of TDP-43^ΔNLS^ C175V/ΔHP inside stress granules in HeLa cells, related to Figure 5

HeLa BAC-G3BP1-mCherry cells with medium expression of TDP-43^ΔNLS^ C175V/ΔHP were stressed by addition of 100 μM arsenite for 60 min and recovery for 120 min. TDP-43^ΔNLS^ C175V/ΔHP (green) and stress granules marked by mCherry-tagged G3BP1 (magenta) were visualized. Scale bar, 10 μm.

Video S7. Intra-condensate demixing of TDP-43^ΔNLS^ inside stress granules in iPS-MN cells, related to Figure 6 iPS-MN cells cotransfected with TDP-43^ΔNLS^ and G3BP1 were stressed by addition of 100 μM arsenite. TDP-43^ΔNLS^ (green) and stress granules marked by mCherry-tagged G3BP1 (magenta) were visualized. Scale bar, 10 μm.

